# Role of RIG-I-like receptors in innate immune sensing of Coxsackievirus B3 and encephalomyocarditis virus in murine macrophages and fibroblasts

**DOI:** 10.1101/441667

**Authors:** Esther Francisco, Mehul Suthar, Michael Gale, Amy B. Rosenfeld, Vincent R. Racaniello

## Abstract

Viral infections are sensed by pattern recognition receptors that trigger an innate immune response through the expression of interferons (IFNs) and other cytokines. Most RNA viruses are sensed by the RIG-I like receptors (RLR)s. The contributions of these receptors to sensing viruses of the *Picornaviridae* family were investigated. Encephalomyocarditis virus (EMCV) and Coxsackievirus B3 (CVB3), picornaviruses of the *Cardiovirus* and *Enterovirus* genera, are detected by both MDA5 and RIG-I in bone marrow derived macrophages. In macrophages from wild type mice, type I IFN is produced early after infection; IFNβ synthesis is reduced in the absence of each sensor, while IFNα production is reduced in the absence of MDA5. EMCV and CVB3 do not replicate in murine macrophages, and their detection is different in murine embryonic fibroblasts (MEFs), in which the viruses replicate to high titers. In MEFs RIG-I was essential for the expression of type I IFNs but contributes to increased yields of CVB3, while MDA5 inhibited CVB3 replication but in an IFN independent manner. These observations demonstrate that innate sensing of similar viruses by RLRs depends upon the cell type.

**Importance:** Enteroviruses such as Coxsackieviruses are the most common human respiratory pathogens. The host’s innate immune response, in particular that modulated by the production of type I and III interferons, is thought to restrict picornavirus infection. Two cytoplasmic proteins, MDA5 and RIG-I, are critical for initiating the early innate immune response against these viruses. Mutations within MDA5 encoding gene have been associated with the development of severe enterovirus associated respiratory illnesses in healthy children. To further understand how the innate immune response dependent upon MDA5 and Rig-I is initiated during picornavirus infection, macrophages from mice lacking MDA5 or RIG-I were infected with Coxsackievirus B3 (CVB3) and a related animal virus. RIG-I is essential for type I IFN production during CVB3 infection; when MDA5 is present, viral titers are reduced by an IFN-independent pathway. These observations demonstrate that innate sensing of viruses by MDA5 and RIG-I depends upon the cell type.

## Introduction

Viruses of the family *Picornaviridae* cause a myriad of important diseases that vary in host tropisms, severity, and pathologies. Coxsackieviruses comprise two groups, A and B, which are responsible for the common childhood hand foot and mouth disease and aseptic meningitis, respectively, and have been implicated in type I diabetes (1). The natural host of encephalomyocarditis virus (EMCV), a member of the *Cardiovirus* genus, is mice although it can infect many other animals and is an important veterinary pathogen that is known to cause outbreaks in farms and zoos, leading to death of animals from encephalitis or myocarditis (2). Poliovirus causes the paralytic disease poliomyelitis in humans and belongs to the *Enterovirus* genus. The diversity in tropisms and diseases caused by viruses with similar genomes makes members of the *Picornaviridae* compelling to study.

A crucial determinant of the outcome of viral infection is the response of the innate immune system, which can limit the ability of viruses to replicate and cause damage at the site of infection and spread to other tissues. Picornavirus infections may be sensed by the cytoplasmic helicase MDA5 (3-10), a member of the RIG-I like receptor protein family (RLR). The other two members of this family are RIG-I and LGP2. These proteins consist of two N-terminal CARD domains, a DExD/H box RNA helicase domain, and in RIG-I a C-terminal repressor domain. LGP2 is similar in structure to RIG-I except that it lacks the N-terminal CARD domains, which may allow it to be a negative or a positive regulator of RIG-I or MDA5 signaling (11-13).

Sensing of viral infection by RLRs begins with the binding of the viral RNA to the carboxy-terminal domain, which leads to ATP hydrolysis and a conformational change in the protein. As a consequence the N-terminal CARDs can interact with the CARD of MAVS, a protein located in the mitochondrial outer membrane. This interaction leads to the activation of IRF3 and NF-κB, whose translocation into the nucleus results in type I IFN expression, and the downstream expression of interferon stimulated genes (ISGs), which create an anti-viral state within a cell (6, 14-18).

We previously showed that both RIG-I and MDA5 are cleaved during infections with multiple picornaviruses (19, 20). These observations led us to determine the roles of RIG-I and MDA5 in the sensing of picornavirus infections. The results of previous *in vivo* experiments in cell cultures and in mice identified MDA5 as a sensor for multiple picornaviruses (reviewed in (21)). However, the presence of multiple cell compartments in animals (22) and strain differences (23) can obscure the involvement of sensors during infection.

By infecting cells from mice lacking the genes encoding MDA5 or RIG-I, we found that both sensors are involved in type I IFN production in response to infection with EMCV and Coxsackievirus B3 (CVB3). MDA5 is essential for type I IFN expression but RIG-I is necessary for maximal levels of IFNβ expression in murine macrophage infections with EMCV and CVB3. Both MDA5 and RIG-I are needed for type I IFN expression after EMCV infection of MEFs. During CVB3 infection RIG-I is essential for type I IFN expression; when MDA5 is present, viral titers are reduced by an IFN-independent pathway. These observations demonstrate that innate sensing of viruses by RLRs depends upon the cell type.

## Materials and Methods

### Cells and viruses

Murine bone marrow from *Mda5*^-/-^ and wild type control mice in a 129×1/SvJ background (from Paula Longhi, Rockefeller University) and *Rig*-I^-/-^ and wild type control mice in a mixed ICR/B6/129 background were were harvested and cultured in DMEM supplemented with 20% L-cell conditioned media, 10% fetal bovine serum (Atlanta Biologicals, Lawrenceville, GA), and 1% penicillin-streptomycin (Invitrogen, Grand Island, NY) for 6-8 days to produce bone marrow derived macrophages.

MEFs were produced from *Mda5*^-/-^ and matched wild type MEFs in a 129×1/SvJ background (from Marco Colonna, Washington University), *Ifnar*^-/-^ MEFs on a 129×1/SvJ background (from David Levy, New York University) and *Rig*-I^-/-^ mice and their matched wild type controls in a mixed ICR/B6/129 background. MEFs were maintained in DMEM supplemented with 15% fetal bovine serum, 1mM sodium pyruvate (Invitrogen), 2mM L-glutamine (Invitrogen), 1% penicillin-streptomycin, 0.1mM non-essential amino acids (Invitrogen).

Viral RNA genomes were produced from plasmids containing DNA copies of EMCV (pEC4) and CVB3 (pSVCVB3) genomes (24, 25) by *in vitro* transcription with T7 RNA polymerase (Fermentas, Glen Burnie, MD). Viral stocks were produced by transfection of transcribed RNA into HeLa cells using DEAE-dextran as previously described (26).

### Virus infections

Infections were performed in 35mm dishes. Cells were counted at the time of infection and viral stocks were diluted in PBS+0.01%BSA (Sigma, St. Louis, MO) for infections at the appropriate MOI. Two hundred microliters of diluted virus were used to infect each plate at 37°C for 45 min with rocking. Cells were then washed twice with PBS and covered with 1mL of appropriate media; samples taken at this time are called 0 time point. Mock infected cells were treated with PBS+0.01%BSA instead of virus.

### Quantitative PCR

Total RNA was harvested using an RNeasy Kit (Qiagen, Valencia, CA) with DNase I treatment (Qiagen) according to the manufacturer’s protocol. Complementary DNA was generated using iScript cDNA synthesis kit (BioRad, Hercules, CA) using both random hexamer and polyT primers according to the manufacturer’s protocol. Complementary DNA was diluted 1:10 and qPCR done on individual genes using Syber Green PCR Master Mix (Applied Biosystems, Foster City, CA) and appropriate primers were used as follows:

IFNβ forward GCACTGGGTGGAATGAGACTATTG,
IFNβ reverse TTCTGAGGCATCAACTGACAGGTC
IFNα forward CCACAGCCCAGAGAGTGACCAG
IFNα reverse AGGCCCTCTTGTTCCCGAGGT
*Mda5* forward GCATGCTGGTCTGCTCGGGA
*Mda5* reverse TCCCCAAGCCTGGCCACACT
RIG-I forward ACTGCCTCAGGTCGTTGGGC
RIG-I reverse GCATCCAGGGCGGCACAGAG
OAS forward CTGCCAGCCTTTGATGTCCT
OAS reverse TGAAGCAGGTAGAGAACTCGCC
β-actin forward GCTGTGCTGTCCCTGTATGCCTCT
β-actin reverse CCTCTCAGCTGTGGTGGTGAAGC
IFNλ2/3 forward AGCTGCAGGTCCAAGAGCG
IFNλ2/3 reverse GGTGGTCAGGGCTGAGTCATT.

Quantitative PCR was performed in triplicate using an ABI 7500 real time PCR machine. Data was analyzed using the 2^ΔΔCt^ method as previously described and expressed as fold change using β-actin as the calibrator gene (27).

### Plaque assays

Samples for virus titration were produced by harvesting cells and medium, followed by three cycles of freezing and thawing and then by centrifugation at 5000xg for 5 min. All titrations were done on monolayers of HeLa cells in six well plates. Ten-fold serial dilutions of virus were made in PBS+0.1 mg/ml BSA (Sigma) and 0.1mL of each dilution was added per well. Plates were incubated at 37°C for 45min with rocking and overlay was then added. For PV and CVB3 a single agar overlay of 0.8% Bacto-agar (Sigma) DMEM, 5%BCS, 0.05%NaHCO_3_, was used. For EMCV a two-overlay system was used. Overlay 1 had a final composition of DMEM, 0.8% Noble agar, 1%BCS, 0.2% NaHCO_3,_ 50mM MgCl_2,_ 1% non-essential amino acids. Overlay 2 had a final composition of DMEM, 0.1%BSA, 40mM MgCl_2_, 0.2% glucose, 2mM sodium pyruvate, 4mM L-glutamine, 4mM oxaloacetic acid (Sigma), and 0.2% NaHCO_3_. PV, CVB3 and EMCV plaque assays were incubated for 48hrs at 37°C. Cells were then fixed with 10%TCA (Sigma) and stained with 0.1% crystal violet (Sigma) in 20% ethanol.

## Results

### RIG-I is important and MDA5 is essential for sensing EMCV in macrophages

To determine if the IFNβ and IFNα responses to EMCV is controlled by either Rig-I and MDA5, we infected bone marrow derived macrophages from *Mda5*^-/-^ mice in a pure 129×1/SvJ background and *rig-I*^-/-^ mice in a mixed background and their matched wild type controls, as done in previous studies (4, 5), and IFN induction was measured by qPCR.

A strong and early induction of IFNβ was observed in wild type cells, peaking at 6 hours post infection in the *Mda5* null background and 3 hours post infection in the *Rig-I* null background (Fig. 1A). To address the differences in the induction of IFN expression between the wild type cells we examined RNA levels encoding the two cytosolic receptors. Cells from both mouse strains express about twice as much *Rig-I* than *Mda5* RNA (Fig. 2). Loss of one of the cytosolic receptors has no effect on the basal level of expression of the other (Fig. 2A). Of note, in wild type macrophages the absolute levels of sensor expression in the mixed background is twice as high as in the 129×1/SvJ background (Fig. 2A). This observation may explain the quicker and sharper induction of IFNβ in macrophages in the mixed background compared to the 129×1/SvJ background.

**Figure 1.**
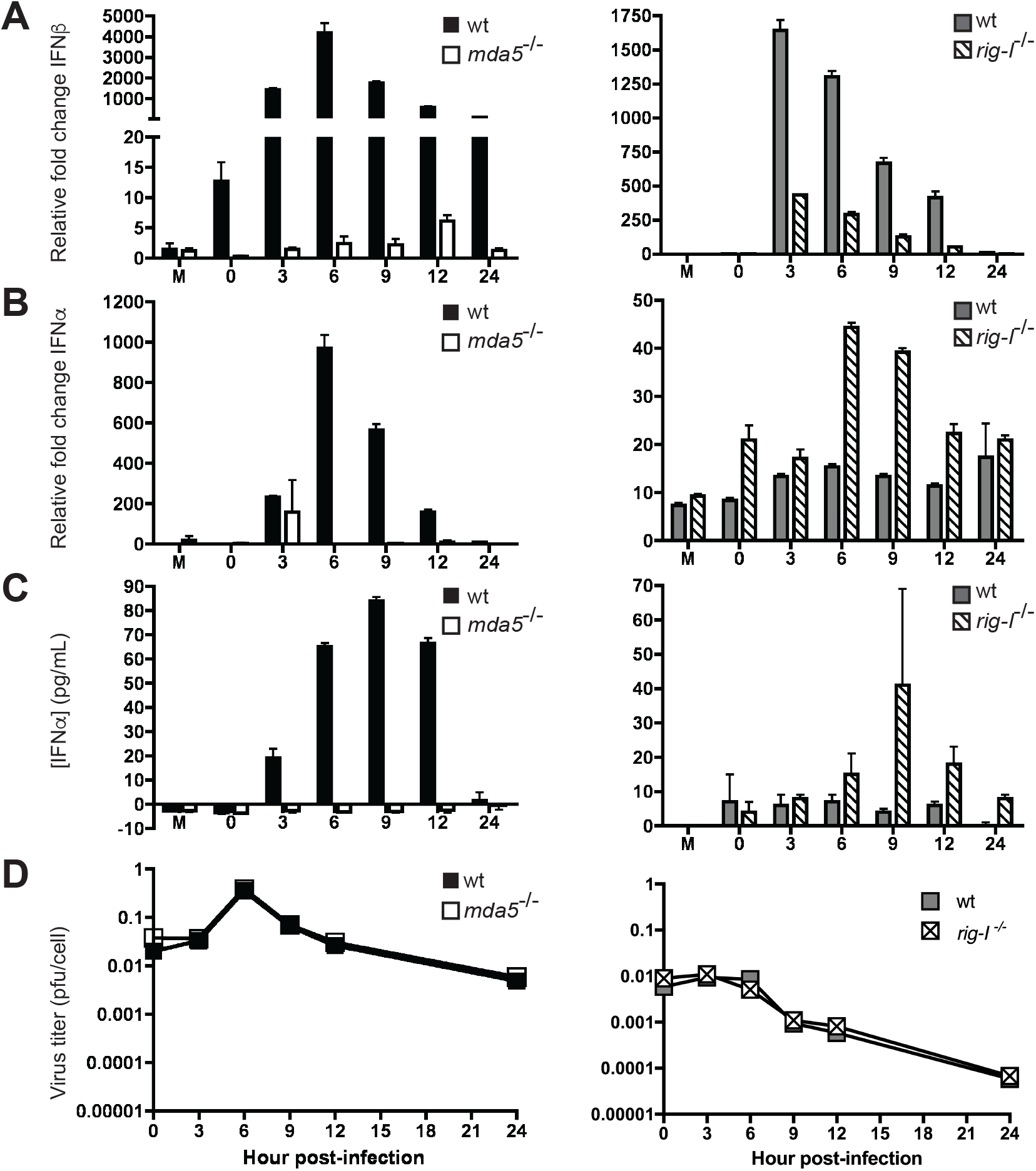
Role of MDA5 and RIG-I in the induction of type I IFN in macrophages infected with EMCV. Bone marrow from *mda5* or *rig-I* null mice and their matched wild type controls were harvested and cultured in conditioned media and infected with EMCV at an MOI=1. Total RNA from mock infected cells and infected cells was harvested at the indicated hours post infection. Expression of IFNβ **(A)** and IFNα **(B)** was determined by qPCR. Supernatants **(C)** were used to measure IFNα protein levels by ELISA. Viral titers during infection were determined by plaque assay **(D)**. Representative data from one of three independent experiments are shown. Plaque assays were performed in duplicate and qPCR performed in triplicate. Error bars show standard deviation within one experiment.

**Figure 2.**
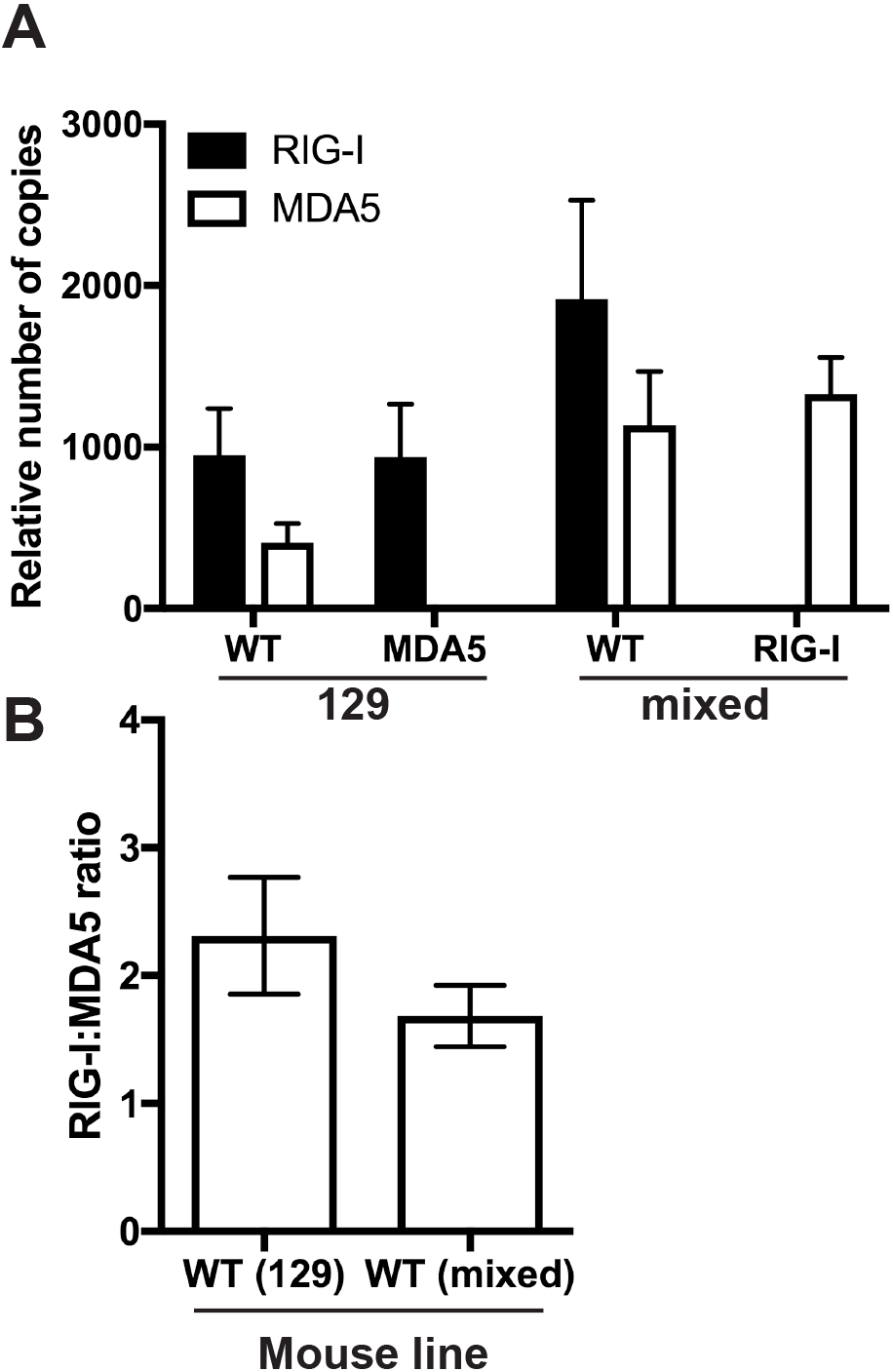
Basal levels of RIG-I and MDA5 expression in macrophages. Total RNA from uninfected bone marrow derived macrophages of different mice was harvested and copies of RIG-I and MDA5 RNA calculated from qPCR using a plasmid standard. **(A)** Copy numbers were normalized to β-Actin Ct values. **(B)** Ratio of relative RIG-I to MDA5 expression was calculated. n=5.

As previously reported, the absence of *Mda5* resulted in a loss of IFNβ expression after EMCV infection (Fig. 1A). Unexpectedly, we found that the absence of RIG-I led to a reduction of IFNβ expression (Fig. 1A). This observation indicates that both MDA5 and RIG-I make contributions to sensing of EMCV infection in murine bone marrow-derived macrophages.

IFNα expression was also induced after infection of both wild type cells, although to a lesser extent (Fig. 1B). As expected, expression of IFNα was not observed in the *Mda5^-/-^* cells. However, in the absence of *rig-I* IFNα was still expressed but at lower levels (Fig. 1B). Similar results were obtained when IFNα protein levels were measured in cell supernatants after infection (Fig. 1C). These observations suggest that in this cell type, sensing of EMCV infection through MDA5 is essential to the expression of IFNβ and IFNα, while sensing through RIG-I only contributes to the expression of IFNβ.

Total virus was harvested at various hours post-infection to monitor the yield of infectious progeny. Yields of infectious EMCV per cell were low in both wild type and *Mda5^-/-^* macrophages (Fig. 1D). Wild type and *rig-I^-/-^* cells did not produce infectious EMCV (Fig. 1D). Absence of RIG-I or MDA5 did not affect virus yields. These observations indicate that the antiviral response is triggered in these cells through MDA5 and RIG-I does not impair viral replication.

### RIG-I and MDA5 sense EMCV in MEFs

To determine the contribution of MDA5 and RIG-I in cells in which the virus could replicate, we infected *rig-I*^-/-^ and *Mda5*^-/-^ MEFs and their wild type controls with EMCV. EMCV replicates well in this cell type (Fig. 3A). Type I IFN was induced in response to infection, peaking at 24 hours post infection (Fig. 3B and C), and induction of the ISG OAS followed (Fig. 3D). The absence of both MDA5 or RIG-I decreased the amounts of IFNβ, IFNα and OAS (Fig. 3B, C and D). These decreases in antiviral response had no effect on viral replication since high amounts of type I IFN were not produced in wild type cells until after the peak of viral replication. However, the decrease in type I IFNs caused by the absence of MDA5 and RIG-I indicate that both proteins are involved in the sensing of EMCV infection in MEFs.

**Figure 3.**
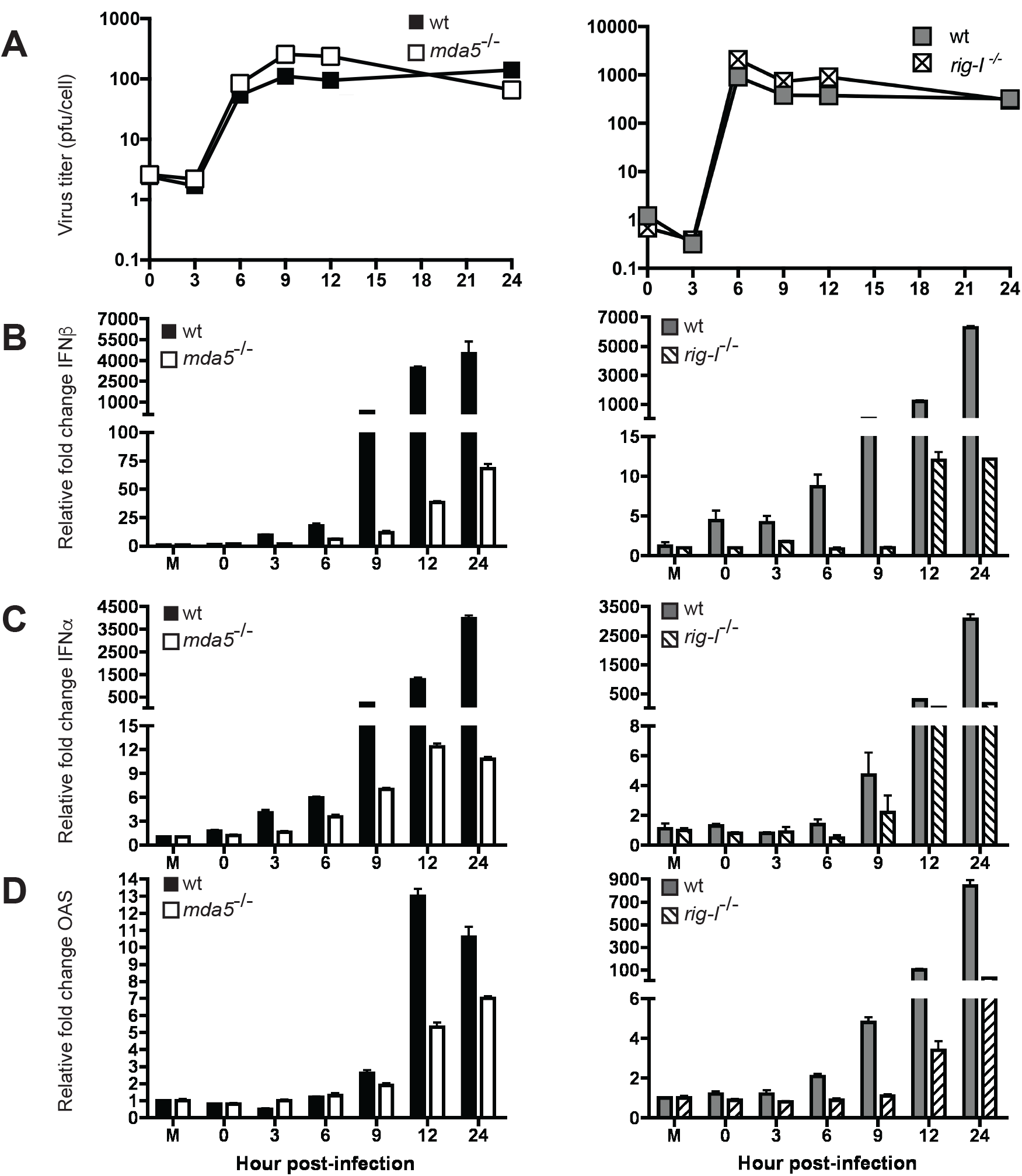
Role of MDA5 and RIG-I in the induction of an antiviral state in MEFs infected with EMCV. *MDA5* and *rig-I* null MEFs and their matched wild type controls were infected with EMCV at an MOI=1. Total virus was harvested at indicated hours post infection and viral titers determined by plaque assay **(A)**. Total RNA was extracted from mock infected cells (M) and infected cells at indicated hours post infection and expression of IFNβ **(B)**, IFNα **(C)**, andOAS **(D)** was determined by qPCR. Representative data from one of three independent experiments are shown. Plaque assays were performed in duplicate and qPCR performed in triplicate. Error bars show standard deviation within one experiment.

To determine whether sensing of EMCV in MEFs is dependent on replication, *Mda5*^-/-^, *rig-I*^-/-^ and matched WT control MEFs were incubated with infectious or UV inactivated virus. Total virus was harvested at 24 hours post infection and plaque assays were performed to ensure that no viral replication occurred in MEFs treated with UV inactivated virus. Replicating virus was sensed in wild type, *Mda5*^-/-^ and *rig-I*^-/-^ MEFs resulting in the induction of IFNβ and IFNα 24 hours post treatment (Fig. 4). UV inactivated EMCV did not induce IFNβ or IFNα in wild type cells, demonstrating that sensing in MEFs requires viral replication (Fig. 4).

**Figure 4.**
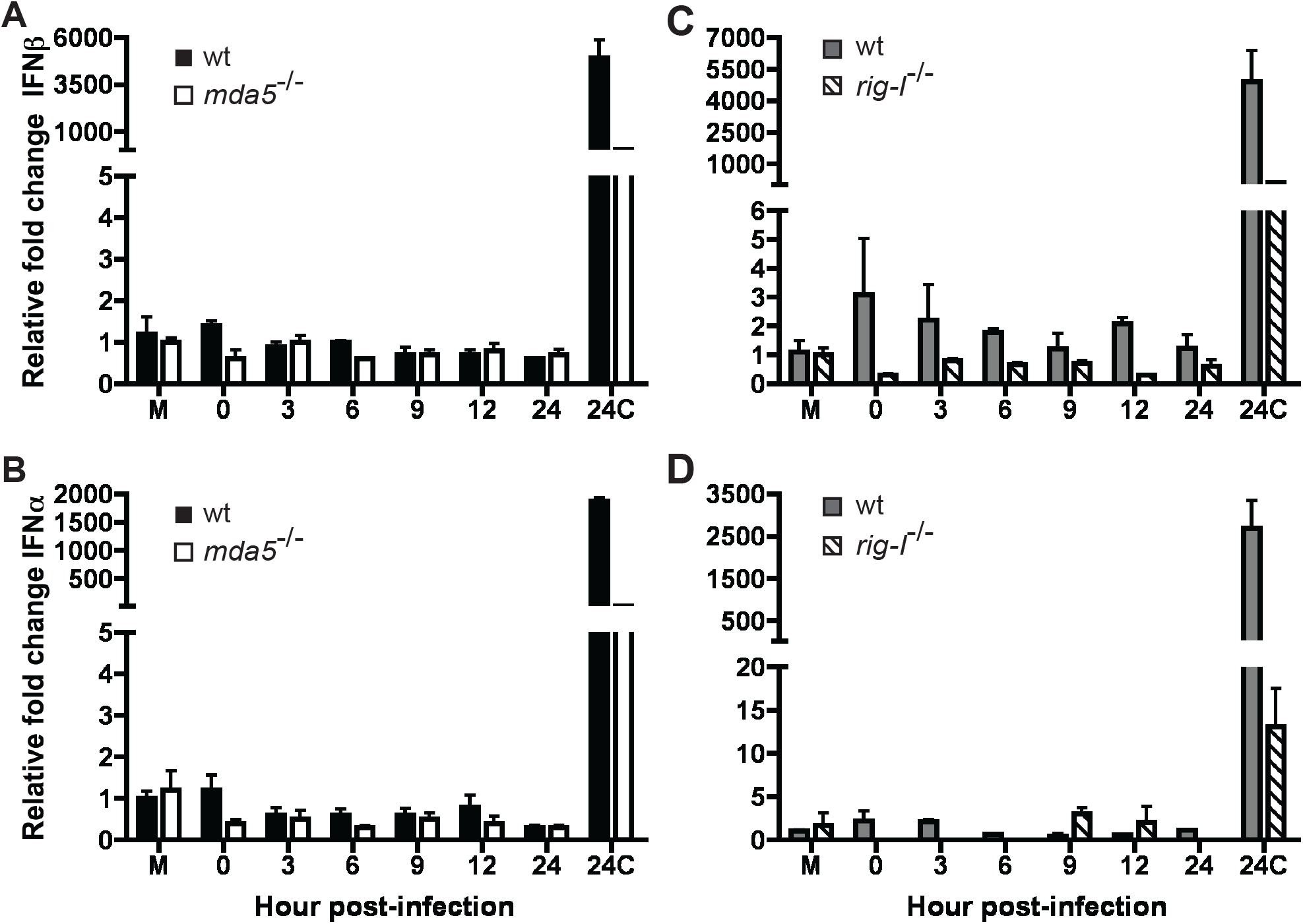
Role of EMCV replication on the induction of an MDA5 and RIG-I dependent antiviral state in MEFs. *MDA5* **(A,B)** and *rig-I* **(C,D)** null MEFs and their matched WT controls were treated with EMCV or UV inactivated EMCV at 1pfu per cell. Total RNA was collected at 24hrs p.i. from mock (M) and EMCV (24C) treated cells as controls and at varying times post UV inactivated EMCV treatment and expression of IFNβ **(A, C)** and IFNα **(B, D)** was determined by qPCR performed in triplicate. Representative data from one of three independent experiments are shown. Error bars show standard deviation within one experiment.

### RIG-I is important but MDA5 is essential for the sensing of CVB3 in macrophages

The role of MDA5 and RIG-I in enterovirus infection was studied using CVB3 infection of *Mda5*^-/-^ and *rig-I*^-/-^ murine cells. Infections were done at high MOI because low MOI did not induce type I IFN, ISGs or cytopathic effect (data not shown). CVB3 infection of bone marrow derived macrophages elicited a type I IFN response, but of a greatly reduced magnitude compared with EMCV (Figs. 1 and 5). The onset of the IFNβ response to CVB3 infection occurred early in the mixed ICR background, peaking at 3 hours post infection, but was delayed in a 129×1/SvJ background, peaking at 12 hours post infection (Fig. 5A). Similar observations were made during infection of macrophages by EMCV (Fig. 1). In the 129×1/SvJ background CVB3 infection resulted in the induction of both IFNβ and IFNα in wild type but not in macrophages lacking MDA5 (Fig. 5A, 5B). In an ICR mixed background only IFNβ is induced in wild type macrophages; in the absence of RIG-I, its levels are reduced. Although IFNα was not induced in WT macrophages of a mixed ICR background, expression was induced in the absence of RIG-I (Fig. 5B). A similar negative regulation of IFNα expression by RIG-I was also observed in EMCV infected macrophages (Fig. 1B). Unlike IFNα expression in macrophages infected with EMCV this negative regulation by RIG-I was not observed in measurements of IFNα protein levels (Fig. 5C). These observed differences in Type I IFN expression are not associated with differences in infectious virus production (Fig. 5D). Similar to our findings with EMCV, a non-productive CVB3 infection of macrophages is sensed by MDA5 to induce an IFNβ and IFNα response. RIG-I contributes positively to the induction of IFNβ but negatively towards the induction of IFNα.

**Figure 5.**
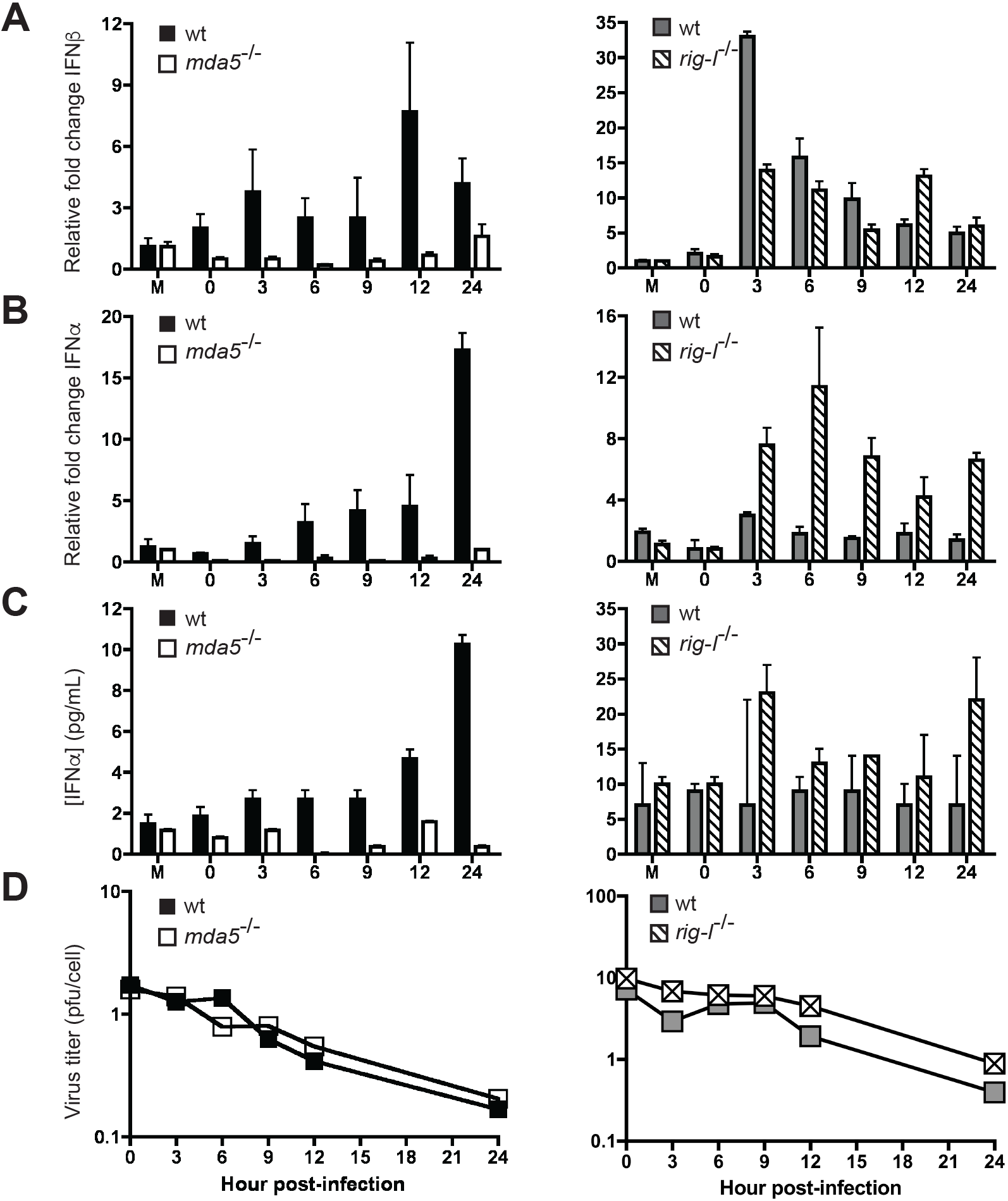
Role of MDA5 and RIG-I in the induction of type I IFN in macrophages infected with CVB3. Bone marrow from *MDA5* or *rig-I* null mice and their matched wild type controls was harvested and cultured in L-cell conditioned media and infected with CVB3 at an MOI=10. Total RNA from mock infected cells (M) and infected cells was harvested at indicated hours post infection. Expression of IFNβ **(A)** and IFNα **(B)** was examined by qPCR. IFNα protein levels were measured by ELISA of culture supernatants **(C)**. Viral titers during infection were determined by plaque assay **(D)**. Representative data from one of three independent experiments are shown. Plaque assays were performed in duplicate and qPCR performed in triplicate. Error bars show standard deviation within one experiment.

### RIG-I is essential for the production of type I IFN during CVB3 infection of MEFs

*Mda5*^-/-^ and *rig-I*^-/-^ MEFs were infected with CVB3 to examine sensing in cells permissive for the production of infectious virus. In WT MEFs, expression of IFNβ, IFNα, and OAS increased in response to infection (Fig. 6). However, in the absence of RIG-I, the expression of IFNβ, IFNα, and OAS was lost (Fig. 6B, C and D), indicating that in MEFs RIG-I is essential for the expression of type I IFN in response to CVB3 infection. However, the loss of type I IFN induction does not correlate with a loss of an effective antiviral state since viral replication decreased in the absence of RIG-I. This observation suggests that the type I IFN response is not protective and that RIG-I may be blocking a protective response (Fig. 6A).

**Figure 6.**
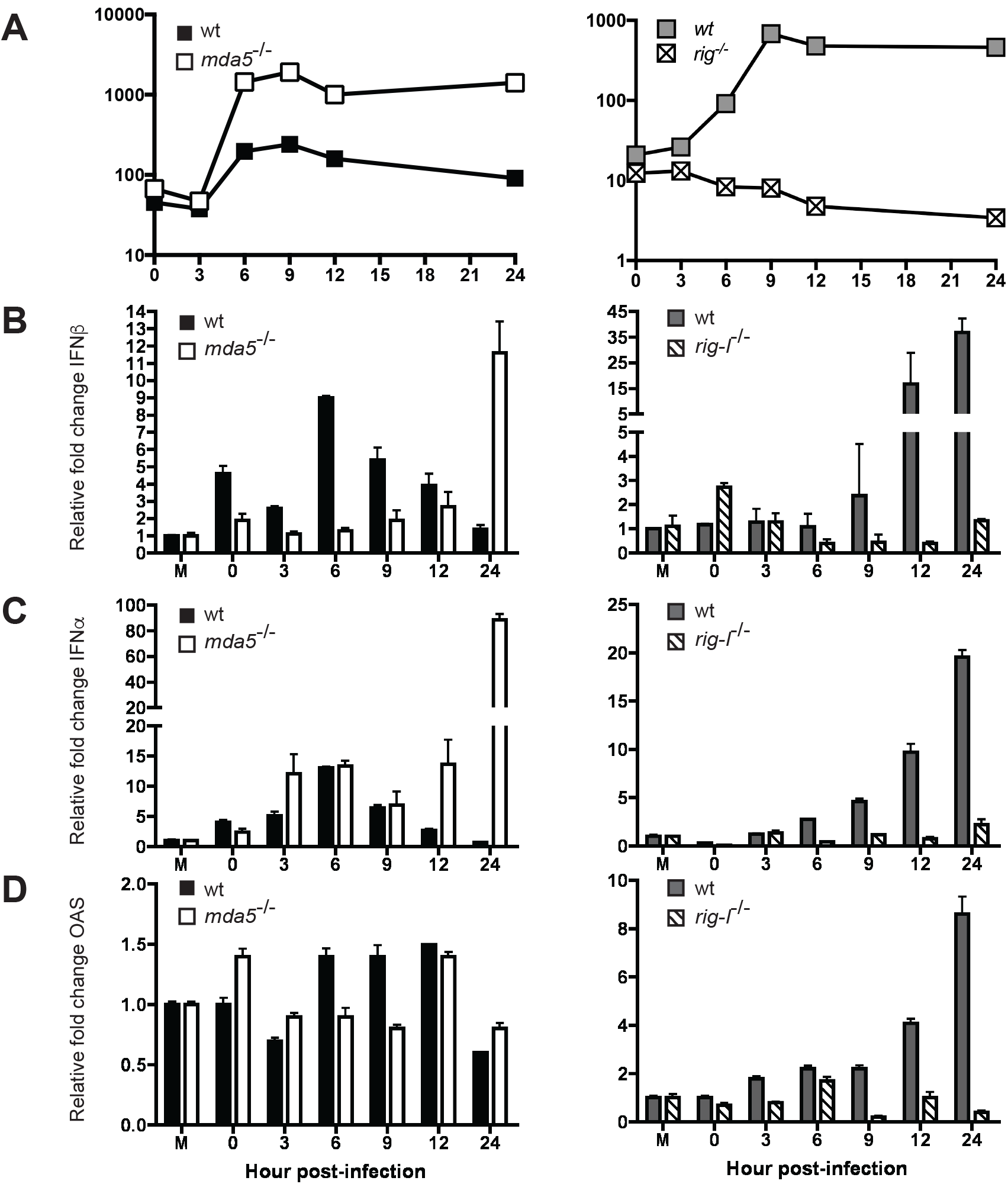
Role of MDA5 and RIG-I in the induction of an antiviral state in MEFs infected with CVB3. *MDA5* and *rig-I* null MEFs and their matched wild type controls were infected with CVB3 at an MOI=50. Total virus was harvested at indicated hours post infection and viral titers determined by plaque assay **(A)**. Total RNA was harvested from mock-infected cells (M) and infected cells at the indicated hours post infection. Expression of IFNβ **(B)**, IFNα **(C)**, and OAS **(D)** was determined by qPCR. Representative data from one experiment of three independent experiments are shown. Plaque assays were performed in duplicate and qPCR performed in triplicate. Error bars show standard deviation within experiment.

Infection of 129×1/SvJ MEFs with CVB3 resulted in a modest increase of IFNβ and IFNα peaking at 6 hours post infection (Fig. 6B and C). In the absence of MDA5 IFNβ and IFNα expression peaked at 24 hours post infection, indicating that MDA5 is essential for the early induction of type I IFNs (Fig. 6B and C). The increase in type I IFN RNA did not, however, lead to an increase in expression of the ISG OAS in either the wild type or the *Mda5*^-/-^ cells (Fig. 6D). Despite the absence of an antiviral state detectable by OAS induction and only late production of type I IFN production in MDA5 deficient MEFs, there is a difference in the replication of CVB3 in the two cell lines. CVB3 replicated to ten fold higher titers in the absence of MDA5 (Fig. 6A), suggesting that MDA5 is important for the induction of an antiviral state.

Treatment of *Mda5*^-/-^, *rig-I*^-/-^ and matched wild type control MEFs with UV inactivated CVB3 showed that the modest amount of type I IFN induced is dependent on viral replication (Fig. 7).

**Figure 7.**
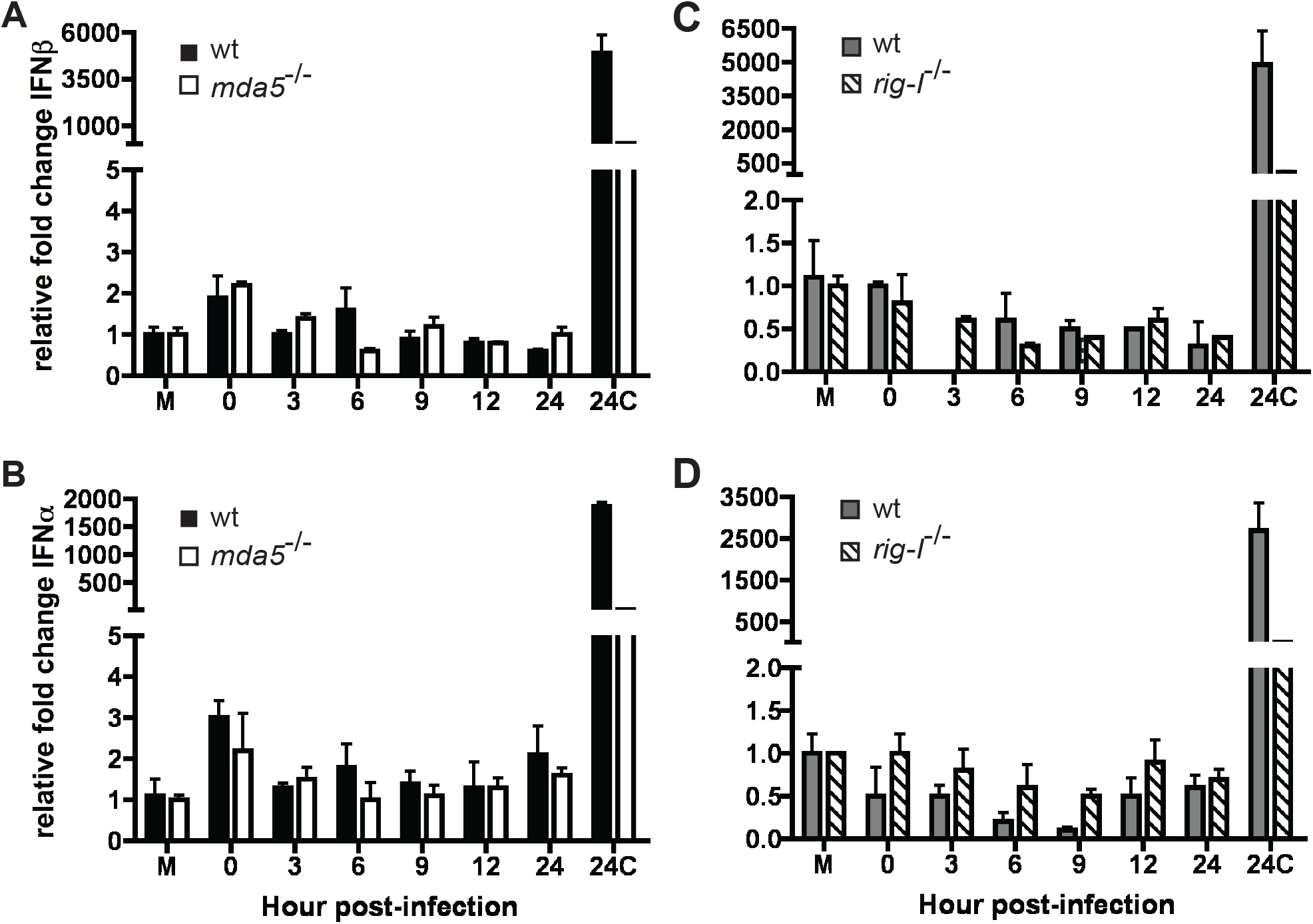
Role of CVB3 replication on the induction of an MDA5 and RIG-I dependent antiviral state in MEFs. *MDA5* **(A, B)** and *rig-I* **(C, D)** null MEFs and their matched WT controls were treated with EMCV or UV inactivated CVB3 at 1pfu per cell. Total RNA was collected at 24 hrs p.i. from mock (M) and EMCV (24C) treated cells as controls and at varying times post UV inactivated CVB3 treatment and expression of IFNβ **(A,C)** and IFNα **(B,D)** determined by qPCR performed in triplicate. Representative data from one of three independent experiments are shown. Error bars show standard deviation within one experiment.

### MDA5 has a role independent of type I IFN in inhibiting CVB3 replication in MEFs

To ensure that the antiviral state produced by MDA5 was independent of type I IFN, we determined CVB3 yields in MEFs unable to respond to type I IFNs due to the knockout of the type I IFN receptor (*ifnar^-/-^*). These MEFs, in a 129×1/SvJ background, were infected with wild type and *Mda5*^-/-^ MEFs and viral titers were determined at different times post infection. As we previously observed, CVB3 titers modestly increased in wild type MEFs, while in the absence of MDA5 viral growth was about ten-fold greater (Fig. 8). This observation was made both at a high MOI of 50 at which type I IFN induction could be detected, as well as at a lower MOI of 1 which did not lead to type I IFN induction. In the absence of IFNAR, viral growth was slightly greater than in wild type MEFs, but not as great as in *Mda5*^-/-^ MEFs (Fig. 8). This observation further suggests that sensing through MDA5 produces an antiviral response that is independent of type I IFNs.

**Figure 8.**
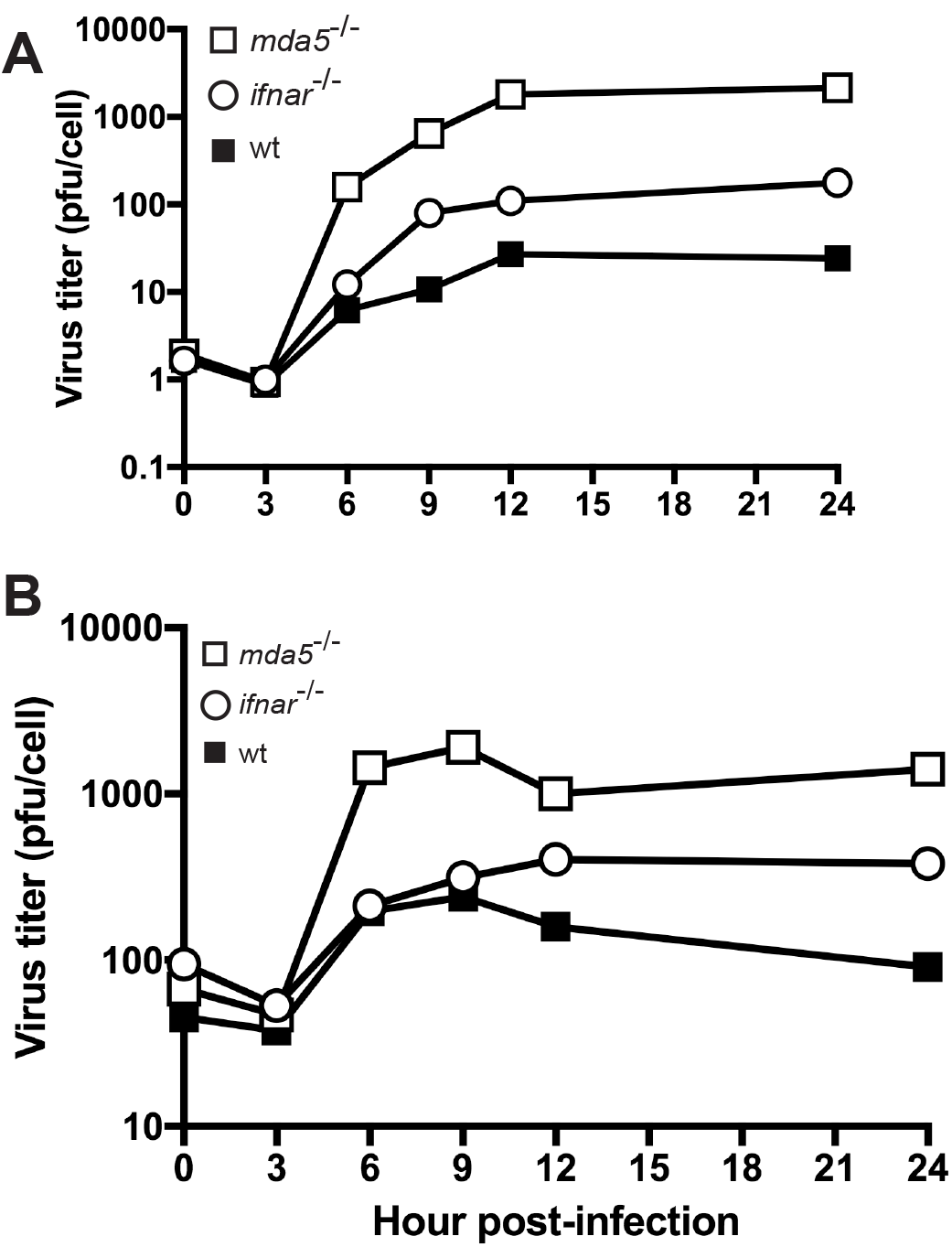
Effect of MDA5 and IFNAR on CVB3 replication. *MDA5* and *ifnar* null MEFs were infected with CVB3 at MOI=1 **(A)** or MOI=50 **(B)**. At various times post infection samples were harvested and total virus titer was determined by plaque assay. Representative data from one experiment of two independent experiments are shown. Plaque assays were performed in duplicate.

Poliovirus but not EMCV can replicate in cells pre-treated with IFNα (28). To determine the effect of IFNα on yields of infectious CVB3, HeLa cells were mock or pre-treated with IFNα at 1000 units/mL, and then infected with CVB3, poliovirus and EMCV. IFNα pre-treatment reduced the levels of CVB3 growth in HeLa cells similar to the extent observed for poliovirus. In contrast, yields of EMCV were substantially reduced by pretreatment of cells with IFNα (Fig. 9).

**Figure 9.**
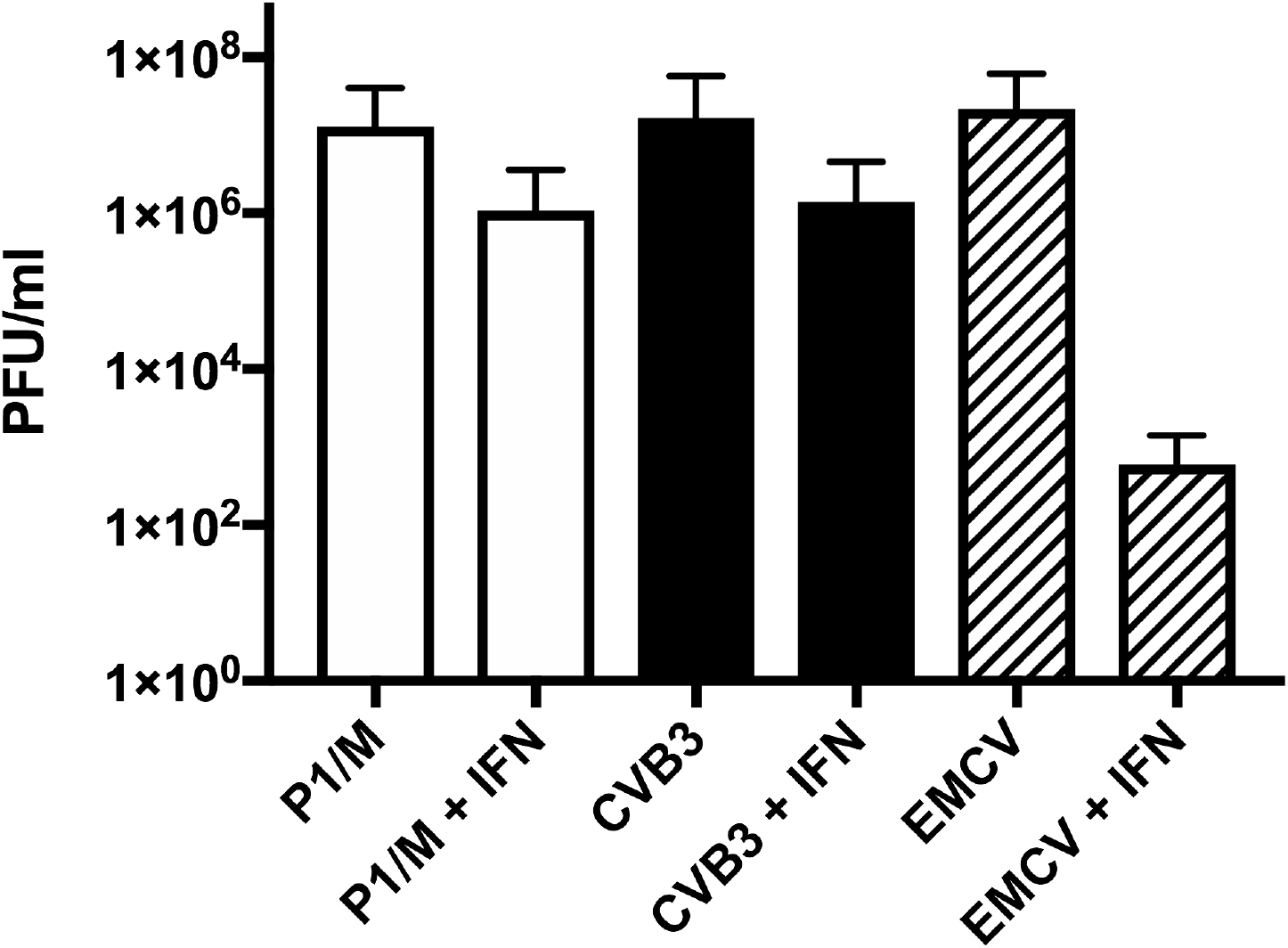
Viral growth in untreated or IFNα pretreated cells. HeLa cells were treated with or without 1000U/mL of IFNα for 16 hours and infected with poliovirus P1/Mahoney (P1/M), Coxsackieviru B3 (CVB3), or encephalomyocarditis virus (EMCV) at an MOI=1. At 24 h post infection total virus was harvested and titers were determined by plaque assay. Results are the average of three independent experiments, and error bars represent standard deviation. Plaque assays were performed in duplicate.

### MDA5 dependent sensing of CVB3 infection of MEFs is independent of type III IFN

We hypothesized that type III IFNs could explain the observed type I IFN independent inhibition of CVB3: absence of MDA5 might result in a loss of type III IFN expression. Type III IFNs use a different receptor for signaling and may preferentially activate a different assortment of STATs to induce a different array of ISGs than type I IFNs. To address this hypothesis, total RNA was collected from wild type, *Mda5*^-/-^ and *ifnar^-/-^* MEFs infected with CVB3. IFNλ expression was not induced in wild type MEFs but was induced in the absence of MDA5 (Fig. 10). The induction of IFNλ2/3 in *Mda5*^-/-^ MEFs was detectable in MEFs infected at the low MOI of 1, at which no induction of type I IFNs was observed (Fig. 10). These findings show that the inhibitory effect of MDA5 on CVB3 replication is independent of both types I and III IFN.

**Figure 10.**
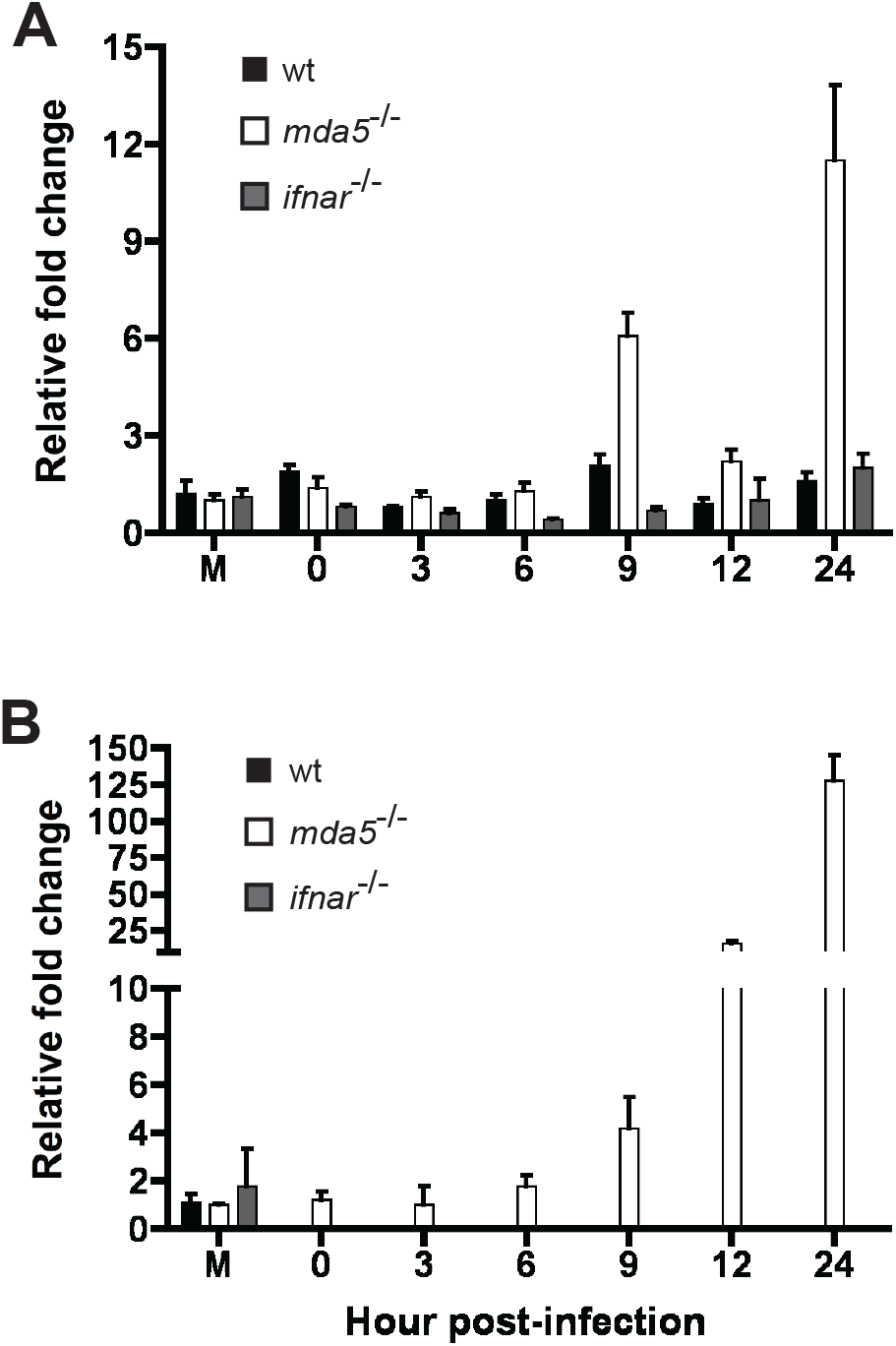
Role if MDA5 and IFNAR on type III IFN expression in MEFs infected with CVB3. Wild type, *MDA5* and *ifnar* null MEFs were infected with CVB3 at MOI=1 **(A)** and MOI=50 (B). Total RNA was harvested and qPCR for murine IFNλ2/3 was performed. Representative data from one of three independent experiments is shown, qPCR performed in triplicate. Error bars show standard deviation within one experiment.

## Discussion

The cytoplasmic receptors RIG-I and MDA5 were initially thought to sense distinct sets of viral RNAs. It is commonly accepted that RIG-I recognizes infections by most families of RNA viruses while MDA5 senses viruses of the *Picornaviridae* family (7, 9, 10, 29, 30). This distinction was made from the results of experiments in which MEFs, bone marrow derived dendritic cells, and mice were infected with an array of different RNA viruses, and type I IFN expression and animal survival was determined. While these initial studies were informative, the results of more recent work have begun to modify the original conclusions. It was believed that RIG-I was the sensor for flaviviruses based on experiments using Japanese encephalitis virus. Infections by the flaviviruses Japanese encephalitis virus, hepatitis C virus, West Nile virus and dengue virus are all sensed by RIG-I, but West Nile virus has also been shown to be sensed by MDA5 and therefore contain structures that are ligands for both sensors (5, 31-33). It has been suggested that picornaviral genomes are not sensed by RIG-I because the 5’- ends of viral genomes lack 5’ phosphates and are linked to VPg. Furthermore, RIG-I is cleaved during infection with poliovirus, echovirus, rhinovirus 16 and EMCV in cells that are pre-treated with IFN (20, 34).

In the study reported here we extend our understanding of how viruses of the picornavirus family are sensed by the innate immune system. CVB3 infection of macrophages deficient in RIG-I resulted in a decreased level of IFNβ expression while infection of RIG-I deficient MEFs resulted in a complete loss of IFNβ expression. These observations suggest that during infection with CVB3, RIG-I is necessary for the maximal induction of IFNβ in macrophages and is essential for the production of type I IFNs in MEFs (Fig. 5 and 6). EMCV infections of RIG-I deficient macrophages and MEFs resulted in decreased levels of IFNβ expression, and in MEFs a decrease in IFNα expression. This observation indicates that for EMCV, RIG-I is necessary for the maximal production of IFNβ in macrophages and for IFNβ and IFNα in MEFs (Fig 1 and 3). Results of infecting RIG-I knockout MEFs and macrophages from RIG-I knockout mice demonstrate that RIG-I does sense picornavirus infections, expanding the number of family members that are detected by both MDA5 and RIG-I.

Previously no work had been done to examine the role of RIG-I in CVB3 infection; however it has been shown that the absence of RIG-I does not lead to a significant effect on type I IFN expression in EMCV infected MEFs and bone-marrow derived murine dendritic cells (5). One explanation for these differences could be due to the use of different cell types: macrophages in our studies, as opposed to dendritic cells in others. Furthermore, our studies in macrophages showed that differences in the induction of type I IFNs were detected early after infection, with levels peaking at three to six hours post-infection in *rig-I*^-/-^ and matched wild type control cells (Fig. 1 and 5) while other studies only reported one time point, at 24 hours after infection. At this time point we found that type I IFN expression was low and levels were comparable between *rig-I*^-/-^ and wild type cells.

Differences between published data and our data in MEFs are likely due to differences in mouse strains. Our *rig-I*^-/-^ MEFs were derived from viable mice obtained by the crossing of heterozygous *rig-I*^-/-^ mice in a mixed 129×1/SvJ and C57BL/6 background with outbred ICR mice. In contrast, other studies used MEFs in a 129Sv X C57BL/6 background in which both *rig-I*^-/-^ and *Mda5*^-/-^ mice were lethal at embryonic day 12.5 (5). Although this background allows for the direct comparison of the roles of MDA5 and RIG-I in infection it cannot be assumed that their roles would be as clearly defined in a viable mouse.

Our finding that RIG-I participates in sensing of picornaviruses indicates that picornavirus infections produce an RNA ligand recognizable by RIG-I. Because the ends of RNA molecules recognized by RIG are thought to be important and the ends of picornavirus genomes are obstructed due to the attachment of a protein, VPg, which serves as a primer for viral RNA replication, how RIG-I detects picornavirus RNA is unknown. After viral RNA has entered a cell, VPg is removed before translation, leaving an unobstructed 5’ monophosphate end which would not be recognized by RIG-I (35- 37). Biochemical studies have demonstrated the ability of RIG-I to recognize double stranded RNA with one strand monophosphorylated, supporting the possibility of RIG-I recognition of picornaviral replicative form (RF) RNA, in which the unlinked genome is bound to a newly synthesized negative strand RNA genome (16). Alternatively, RIG-I might sense an unprotected 5’ triphosphate present on the nascent RF RNA (38). Because the levels of RF RNA within a cell are low, it is also possible that RIG-I is sensing a more abundant RNA structure such as the IRES. This highly structured RNA domain contains many double stranded regions, which may be ligands for RIG-I. Because the true definition of a RIG-I ligand remains controversial, further studies are needed to determine how RIG-I recognizes picornavirus RNAs.

EMCV or CVB3 infection of macrophages lacking MDA5 or RIG-I results in a decreased expression of type I IFN but does not result in an increase in viral growth since both EMCV and CVB3 replicate poorly in macrophages (Fig. 1 and 5). Although both viruses do replicate well in MEFs, the absence of RIG-I or MDA5 has no effect on EMCV replication despite the repression of type I IFNs and ISGs (Fig. 3).

The absence of RIG-I during CVB3 infection however appeared to inhibit the formation of an antiviral state by preventing the induction of type I IFN and the ISG OAS. Interestingly there was no concomitant increase in viral replication compared with wild type cells; rather a decrease in viral replication was observed (Fig. 6). The absence of MDA5 did not affect induction of type I IFNs during CVB3 infection; however, an increase in viral titers was observed (Fig, 6A). This inhibitory effect of MDA5 on CVB3 replication was apparent not only at the high MOI of 50 which was required to observe induction of type I IFN, but also at a lower MOI of 1 at which no increase in type I IFNs could be detected (Fig. 8). The increase in viral titer in the absence of MDA5, together with our observed decrease in viral titers when RIG-I is not present, implies that there must be factors independent of type I IFN expression that are inhibiting CVB3 replication. This conclusion is consistent with the results of infection of *MDA5*^-/-^ mice where no significant increase in IFNα serum levels or IFNβ or OAS expression in tissues was detected, yet there was a transient early increase in CVB3 titers, which led to increased tissue damage and disease (23). The results of infecting *ifnar^-/-^* MEFs confirmed this type I IFN independent inhibition of CVB3 replication, since viral titers in these cells were lower than those in *MDA5^-/-^* MEFs at both an MOI of 1 and an MOI of 50 (Fig 8). Viral titers in *ifnar^-/-^* MEFs were slightly higher than in wild type MEFs, indicating that type I IFN does have a slight role in inhibiting CVB3 replication but is not fully responsible for the MDA5 dependent inhibition of CVB3 replication. Additionally, CVB3 was found to be able to replicate in the presence of an antiviral state induced by type I IFN, further supporting the idea that the MDA5 response is independent of type I IFN (Fig. 9).

The poliovirus 2A^pro^ protease determines whether infectious virus is produced in the presence of IFN, in particular a tyrosine at residue 88 of 2A^pro^ which may be involved in the discrimination of substrates (28, 39). The Coxsackievirus 2A^pro^ protease has a tyrosine at the corresponding position, suggesting that the ability of CVB3 to replicate in the presence of type I IFN is a consequence of 2A^pro^ mediated antagonism of inhibitory IFN-stimulated gene products.

Type III IFNs have been shown to be under the control of the transcription factors NFκB, AP-1 and IRFs similar to type I IFNs (40). Therefore, types I and III IFNs are frequently induced in response to the same triggers, although differences in the expression of the two IFNs have been noted. For example, macrophages infected with herpes simplex virus produce type I IFN but not type III IFN, and epithelial cells infected with influenza A virus produce higher levels of type III IFNs than type I IFN (41). These differences are likely due to the dependence of each IFN on various transcription factors: type I IFNs are strongly dependent on IRFs while type III IFNs are more dependent on NFκB (42, 43). There are also some differences in the signaling of type I and III IFNs since the former produce a positive feedback loop in which they up-regulate the expression of more type I and type III IFN. Type III IFN signaling however does not result in an increase in type I or type III IFN expression (42). We hypothesized that type III IFNs could explain the observed type I IFN independent inhibition of CVB3: absence of MDA5 might result in a loss of type III IFN expression. However, we found that expression of IFNλ2/3 was not decreased, but rather induced, in the absence of MDA5. Induction of these cytokines occurred late in CVB3 infection, similar to the kinetics of type I IFNs, beginning at nine hours post infection, and therefore is not likely to not have a causative role in the increase in viral replication in MDA5 deficient MEFs, which commences earlier in infection (Fig. 10).

It is known that viral sensing results in the induction of many cytokines under the control of NFκB in addition to IFNs. One or more of these may be contributing to decreases in CVB3 replication. A more appealing explanation is that MDA5 is signaling through MAVS on the peroxisome. It has been found that an IFN independent antiviral state may be induced by RIG-I through signaling using peroxisomal MAVS rather than mitochondrial MAVS (44). This signaling was found to induce expression of ISGs directly through the activation of IRF1 and to do so before the induction of type I IFNs. If MDA5 can also signal through peroxisomal MAVS, it could induce an effective initial antiviral response that results in a decrease in CVB3 titers as we observed early in infection. Alternatively, activation of IRF3 by MDA5 may allow for transcription of a subset of antiviral ISGs independent of autocrine/paracrine signaling (45).

If RIG-I signaling in this context only occurs through mitochondrial MAVS it might also explain why the induction of type I IFNs and ISGs by RIG-I does not limit viral replication but promotes it. Type I IFN signaling does not affect viral replication because it occurs well after viral replication is complete. Furthermore, the sensing of viral RNA by RIG-I may compete with the ability for MDA5 to carry out the same function. Therefore, in the presence of RIG-I the early, effective, peroxisomal response through MDA5 could be dampened and viral replication is then greater than in the absence of RIG-I. Further experiments are needed to determine how MDA5 is limiting viral replication independent of interferons.

These findings demonstrate the involvement of RIG-I in the sensing of picornavirus infections and the ability of MDA5 to inhibit viral replication in an IFN-independent manner. They also emphasize that the virus, cell type, and genetic background of mice can influence how different viruses are detected by the innate immune system.

## Acknowledgments

This work was supported in part by Public Health Service Grants AI50754, AI139775, AI118916, and AI127463 from the National Institute of Allergy and Infectious Diseases.

We thank Ann Palmenberg and Nora Chapman for EMCV and CVB3 DNA clones respectively; Marco Colonna, David Levy, and Paula Longhi for Mda5^-/-^ MEFs, IFNAR^-/-^ MEFs and MDA5^-/-^ mice.

